# Fibrinogen and complement factor H: The two crucial markers of Parkinson’s disease with cognitive impairment

**DOI:** 10.1101/2023.10.25.563920

**Authors:** Aditi Naskar, M Sachin, Senjuti Sengupta, Pallavi Bhadrachalam, Shantala Hegde, Ravi Yadav, Pramod Kumar Pal, Phalguni Anand Alladi

## Abstract

Animal model-assisted validation is vital for biomarker studies. It provides better means for understanding disease pathogenesis and opens avenues for addressing therapeutics. Our earlier non-targeted label free proteomics-based biomarker study on CSF of Parkinson’s disease (PD) patients with cognitive impairment (PDCI), revealed the presence of elevated levels of fibrinogen and complement factor H (CFAH) in PDCI-CSF. In the present study, we aimed to determine if these proteins harbor a pathogenic potential, when present above physiological levels. Native fibrinogen and recombinant CFAH were intraperitoneally injected in adult C57BL/6J mice and 48h later, motor and cognitive behavioral deficits as well as their neuroanatomical correlates were noted. The reduction in stride length in fibrinogen and CFAH injected mice indicate shuffling gait, often seen in PD patients. Motor deficit was complemented by the loss of dopaminergic (DA) neurons in SNpc, compensatory hypertrophy of the surviving neurons, and reduction in TH expression in striatum, thus reminiscing PD pathology. The low discrimination index in (novel object recognition) NOR test complemented by morphological alterations in Nissl-stained CA-1 and subiculum neurons in the fibrinogen-injected mice imply recognition memory deficits. Thus, our study provides objective evidence that fibrinogen and CFAH harbor the potential to induce motor deficits and cognitive impairments in mice, akin to the PDCI-associated neuro-pathological deficits, and thus are potential biomarkers.

## Introduction

Parkinson’s disease (PD) or the shaking palsy, is quintessentially characterized by a discernible loss of dopaminergic (DA) neurons in the substantia nigra pars compacta (SNpc) in a caudo-rostral and latero-medial fashion (Damier et al., 1999) accompanied by presence of Lewy bodies (reviewed by Goedert et al., 2017). The latter are enriched with α-synuclein, alongside many other proteins (Spillantini et al., 1997; Yang et al., 2022). Primarily, this disorder is identified by motor symptoms like tremor, rigidity, bradykinesia and postural instability (Fahn 2003). In recent times, besides the traditional importance of motor symptoms, the association of non-motor symptoms with PD is gaining importance. Cognitive impairment (CI) is one such non-motor symptom which is common in PD patients and ranges from mild form (Aarsland et al., 2010) to severe dementia (Hely et al., 2008). Hitherto, the final confirmation of PDCI in patients have solely relied on autopsy-based investigations. Introducing a biomarker to the investigations may mitigate this limitation by facilitating the diagnosis, prognosis, and even early prediction of cognitive impairment associated with PD enhancing better management of the disease. A few PET imaging studies showed accumulation of tau in hippocampus and cerebral cortex of PD patients with cognitive impairment (Buongiorno et al., 2017, Hansen et al., 2017). However, in patients worldwide as well as in Asian Indians, there is a paucity of studies to understand the neuro-pathological basis of cognitive deficits in PD. Under these circumstances detailed studies in animal models may serve the purpose better, which are thus far rare. We therefore hypothesize that validation of identified biomarkers using animal model can serve a dual purpose, one of providing an animal model for in-depth studies on disease mechanisms and secondly, facilitating the identification of the biomarker, that is likely to be more potent.

Based on our previous non-targeted LC-MS/MS followed by ELISA-based confirmations we found two proteins viz. fibrinogen and complement factor H (CFAH/CFH), to be greatly upregulated in PDCI. For example, the fold change of different fibrinogen chains in CSF of PDCI patients were FGG (9.2), FGB (7.4), FGA (4.5) (Naskar et al., 2022) and validation result by ELISA showed an average FGG concentration of 513 ng/μl in the CSF. Earlier independent studies on knockout mice, partly explained the role of fibrinogen and CFAH in neurodegeneration, albeit not in PD. For example, Akassoglou et al., (2002) showed accelerated myelination following sciatic nerve injury in mice lacking fibrinogen-α chain. Cortes-Canteli et al., (2010) demonstrated that in AD transgenic mice CRND8, genetic reduction of fibrinogen-α chain (fgα+/−) alleviates cerebral amyloid angiopathy (CAA) and cognitive impairment. Another set of studies provided evidence of physiological role of CFAH. Kam et al., (2016) observed retinal pathology such as retinal β-amyloid deposition in aged cfh−/− mice, suggesting that factor ‘h’ is important for retinal development as also demonstrated in cfh−/− mice by Sivapathasuntharam et al., (2019).

Fibrinogen is a dimeric protein, with each dimer subunit consisting of three non-identical peptide chains termed as Aα, Bβ and γ and it plays a pivotal role in blood clotting. Another protein plasmin assists the lysis of these blood clots (Bardehle et al., 2015). An imbalance between coagulation and fibrinolysis leads to disease pathology (Petersen et al., 2018). Blood-brain barrier (BBB) breach is a common phenomenon, which is concatenated with normal aging and almost all neurodegenerative disorders like Alzheimer’s disease (AD), Parkinson’s disease (PD) etc. Following a BBB breach, perivascular tissue factors and infiltrated proteins convert fibrinogen into fibrin. The neurons and glia are sources of prothrombin and thrombin that aid the conversion (Arai et al., 2006). A couple of studies have reported enhanced levels of fibrinogen in the brain parenchyma of post-mortem AD brains. Fibrin deposition found in transgenic mice models (Paul et al., 2007) exacerbated AD pathology and amyloid-β pathology accentuates fibrinogen deposition in AD patients (Cortes-Canteli et al., 2010).

The complement system is the preliminary line of defense in the humoral immune system. CFAH, an abundant serum complement regulator glycoprotein, inhibits over activation of the alternative pathway (Schwaeble et al., 1987). Increase in the CFAH levels in serum and CSF could be a compensatory mechanism to combat over activation of the alternative pathway in a progressive and relapsing disorder like multiple sclerosis (Ingram et al., 2010). Both fibrinogen and CFAH are indicators of neuroinflammation, one of the major causative factors of neurodegeneration in PD. Neuroinflammation is indicated by elevated levels of pro-inflammatory cytokines viz. TNF-α, interleukin (IL)-1β, IL-6 in the striatum and nigra of both experimental mice and brains of PD patient (Abhilash et al., 2022 preprint; Mogi et al., 1994; Blum-Degen et al., 1995; Müller et al., 1998). In the current study, we upregulated the levels of fibrinogen and CFAH in C57BL/ 6J mice by intraperitoneal injections and investigated whether, they are individually capable of causing pathogenicity in the regions like substantia nigra, striatum, CA-1 and subiculum that are associated with PD as well as PDCI.

The different segments of mouse colon are distinguishable based on the appearance of the mucosa. The caecum and colonic mucosa has deep crypts, but no villi; oriented perpendicular to the long axis of the colon. The crypts are lined by simple columnar absorptive colonocytes, goblet cells, and enteroendocrine cells (Barker et al., 2008). The tunica muscularis has inner circular and outer longitudinal layers. Gut-associated lymphoid tissue (GALT) is abundant in mice and is usually located in the anti-mesenteric submucosa. Large organized areas of lymphocytes known as Peyer’s patches are found to be more prominent in young mice (Jung et al., 2010). Smaller aggregates of lymphoid cells known as cryptopatches or isolated lymphoid follicles can be found along the length of the intestine (Pabst *et al*. 2005) and act as local sites for production and maturation of γδ and αβ T lymphocytes, independent of the thymus gland. Cryptopatches are separate from Peyer’s patches (Ishikawa et al., 1999)

PD affects neurons outside the brain and interferes with normal communication between CNS, oesophagus and stomach, while affecting the enteric nervous system (ENS). PD affects the peristaltic action of intestinal muscles, slackening their movement and resulting in constipation. Pathogens are deemed to enter through olfactory or GI system, leading to Vagus nerve-mediated spread of Lewy bodies to CNS (Braak et al., 2003), and thereby leading to development of constipation and anosmia as prodromal non-motor symptoms.

Gait impairment is one of the hallmark features of PD, hence to ascertain the motor deficits in C57BL/6J mice we determined a spatial gait parameter, following fibrinogen or CFAH administrations. In order to avoid the complexity of other cognitive tasks that entail long training periods as well as deprivation of food or water; we selected a simpler task i.e., novel object recognition (NOR) to assess cognitive deficit, based on recognition memory (Magen et al., 2012). In addition to determining the pertinent behavioral deficits we performed immunohistochemistry-based evaluation of tyrosine hydroxylase (TH) in the substantia nigra and striatum of C57BL/6J mice as a parameter of neuropathological aspects of the motor deficit. We used the optical fractionator probe of the unbiased stereology to quantify the number of nigral DA neurons, nucleator probe to estimate overall neuronal size as well as densitometry to evaluate the cellular expression of TH in the SNpc and the DA inputs to the striatum. The neuropathological characteristic of the cognitive deficit was studied by assessing the volume of two subfields of the hippocampus viz. CA-1 and subiculum with the help of the Cavalieri estimator as well as the cytomorphology of the nucleus and cytoplasm of the neurons of these subfields by morphometry using LAS-X software (Leica Microsystems, Germany). In addition, the gut microarchitecture was also studied following intraperitoneal injection of fibrinogen and CFAH.

## Materials and methods

### Experimental animals

Ethics-related permissions were obtained from the institutional animal ethics committee (AEC/64/382/N.P.) and the experiments were conducted according to the CPCSEA and NIH guidelines. Briefly, 15-17 weeks old C57BL/ 6J mice (Vidyadhara et al., 2019, 2021) were maintained in the Central Animal Research facility (CARF), NIMHANS. We used both female and male mice weighing approximately 20g and 25 g respectively. A 12/ 12h light dark cycle was maintained while housing the animals and they were provided with ad libitum food and water.

### Intraperitoneal injection

Our earlier findings on the pole test (Naskar et al 2022), assisted us in restricting our investigation to two different concentrations viz. 2.5 μg and 5μg of native fibrinogen (ab233609 AbCam UK) and recombinant CFAH (4999-FH-050 R & D systems) in this study. PBS was used to reconstitute the proteins to make it to a total volume of 200 μl for the i.p. injections in C57BL/ 6J mice. The mice were segregated into different experimental groups, 48h post injection, for neurobehavioral and neuropathological studies. The study groups were labeled as: vehicle, fibrinogen 2.5/Fib-2.5, fibrinogen 5/Fib-5, CFAH-2.5, and CFAH-5. The data was obtained from 5 to 7 mice per experiment, encompassing various concentrations or study groups, and processed for statistical significance.

### Gait analysis

Stride length, one of the many primary spatial gait parameters was measured by the Catwalk XT 10.6 gait analysis system (Noldus Information Technology, Netherlands) (Walter et al., 2020). The catwalk system consists of an enclosed glass walkway that is lit up by green LED light and the ceiling by a red LED light. It measures the linear distance between two successive contacts of the same paw, captured by a high-speed color camera connected at the bottom of the walkway and digital signal analysis was done by the inbuilt CatWalk XT 10.6 software. Each mouse was habituated to walk on the walkway (40 cm) for 7 consecutive days. Following 7 days of habituation, a gap of 48 hours was provided before the test session. A threshold of 0.1 and a camera gain of 30 was set to detect all the experimental parameters. A compliant run was assigned to any run between 0.5s and 9.00s. An average of 4 compliant runs were recorded before and 48h after the injection. The runs were classified and gathered for all four paws.

### Novel object recognition (NOR) test

This a behavioral test to assess cognitive abilities/ recognition memory in mice (Ennaceur, 2010). In the original definition of NOR, by Ennaceur & Delacour (1988) an exploration is defined as “orienting the nose towards the object within a distance of 2cm”. Briefly, during the habituation phase each individual mouse was allowed to explore a wooden box of 36 × 36 × 36 cm dimensions for 5m. Twenty-four hour post habituation, each mouse was familiarized with two identical objects placed in diagonally opposite positions, in the open field arena. The test session was conducted 24h post familiarization, in which one of the familiar objects was supplanted by a novel object and the mouse was again allowed to explore for 5m (Leger et al., 2013). Object discrimination index and percentage of exploration were noted.

### Tissue processing for immunohistochemistry

C57BL/6J mice were perfused transcardially using 0.9 % saline followed by 4% paraformaldehyde (0.1M phosphate buffer pH 7.4) as per our established protocol (Vidyadhara et al., 2017; Bhaduri et al., 2018; Seshadri & Alladi 2019). The harvested brains were post-fixed in the same buffer for 48h, at 4°C. The cryoprotected brains were sectioned into 40 μm thick cryosections of SNpc and striatum, which were collected on gelatin subbed slides and were further subjected to TH immunohistochemistry (Vidyadhara et al., 2017; Bhaduri et al., 2018). Briefly, following permeabilization with PBS-Tx, the endogenous peroxidase activity was quenched with 0.1% H_2_O_2_, prepared in 70% methanol for 30 mins and blocked with 3% buffered BSA for 4h at room temperature (Vidyadhara et al., 2017). Thereafter, the sections were incubated with rabbit-polyclonal anti-TH antibody (1:500, ab112 AbCam, UK) for 72h at 4°C. Subsequently the sections were incubated with anti-rabbit secondary antibody (1:200 dilution; 4°C; 17h.; Vector Laboratories, Burlingame, USA) followed by tertiary labelling with avidin biotin complex (1:200, Vector Laboratories; USA; 4h at room temperature). We used 3’-3’-diaminobenzidine dissolved in acetate imidazole buffer (0.06%; 3mg/5ml, 0.1 M, pH 7.4); supplemented with 0.1% H_2_O_2_ to develop the stain.

### Stereological quantification

In order to count the TH-ir cells, every sixth SNpc section was delineated (Fu et al., 2012; Vidyadhara et al., 2017) under the 10X objective of Olympus BX51 Microscope (Olympus Microscopes, Japan) furnished with the Stereo-Investigator (version 2020.1.2.; Micro-brightfield Inc., Colchester, USA), using the optical fractionator probe (Vidyadhara et al., 2017).We quantified the area of the DA neurons simultaneously, with the help of the nucleator probe using the 100X objective. A grid interval of 150 μm on X&Y axes and a square counting frame of 60 μm length were applied (Vidyadhara et al., 2017). The average mounted thickness was set at 25±0.2 µm and a guard zone of 4μm on either side resulted into a depth of 17 µm in the z-direction. While using serial sections, both the hemispheres commencing from the rostral end to the caudal edge, were considered. The numbers of neurons from both the hemispheres were pooled. The volumetric estimation of the striatum was performed by the “Cavalieri estimator” probe of the same software. We earmarked the boundaries of the dorsal and ventral striatum using the 10X objective of a bright-field microscope (Olympus BX51 Microscope Japan). As per the formula, the selected region of interest accounts for the “area” and the total volume is assessed by the total number of sections analyzed.

### Densitometry-based image analysis

The relative amount of protein in the neurons of the SNpc was quantified by densitometry-based image analysis. 8-bit images of the nigral neurons were captured using the Leica ICC50W microscope attached to a digital camera. Images of at least 200 TH immunoreactive DA cells per animal were quantified using the in-built LAS-X software. In case of the striatum, low magnification (4X) images were captured and regions of interest (ROI) were earmarked using the “polyline tool”. Following quantification of the cells and ROIs, gray values and “area” dimensions were used to compute the cumulative values.

### Hematoxylin staining

We studied the cellular morphology of neurons of the hippocampal subfields CA1 and subiculum using hematoxylin stained sections. The same slides were used to assess the area of the ventricles. Similarly, we examined the H&E stained colon sections to assess alterations in the crypt length, number of folds and thickness of the *Tunica muscularis*. Briefly, equilibration of the frozen sections was performed in phosphate buffer for 10m. Thereafter, the sections were incubated in Harris haematoxylin for 45-70s, followed by thorough wash under running tap water followed by lithium exposure (bluing agent) for 30s. The sections were dehydrated in increasing grades of ethanol i.e. (70%, 90% and 100%) and cleared in xylene. Finally, the sections were mounted with dibutyl phthalate polystyrene xylene (DPX).

### Cyto-morphometry

The cyto-morphometric assessments of the CA1, subicular neurons and colon were performed on images captured with Leica ICC50W microscope (Leica Microsystems, Germany). Serial images were captured from four sections each of CA1 and subiculum from each hemisphere and were quantified using the LAS-X software. The “polyline” tool was used to earmark the cytoplasm and the nucleus. At least 15 profiles of cytoplasm and nucleus from each image, i.e., 150-200 cells/specimen were thus analysed. The alteration in gut microarchitecture was analysed on the 4X and 10 X magnification images, using length based evaluation protocols (n=3-5 mice per vehicle/controls and experimental protocol).

## Results

### Fibrinogen and CFAH induce distinct behavioral deficits in mice

#### Both fibrinogen and CFAH affect stride length

Gait analysis revealed an important motor deficit i.e., shortening of the stride length (Figure 1B) in right front foot (RF) of fibrinogen-injected (control vs Fib 2.5 **p= 0.0058) and CFAH-5 injected (control vs CFAH 5 ***p= 0.0009) groups as well as the left front foot (LF) of CFAH injected mice (control vs CFAH 5 *p= 0.0489).

**Figure 1.**
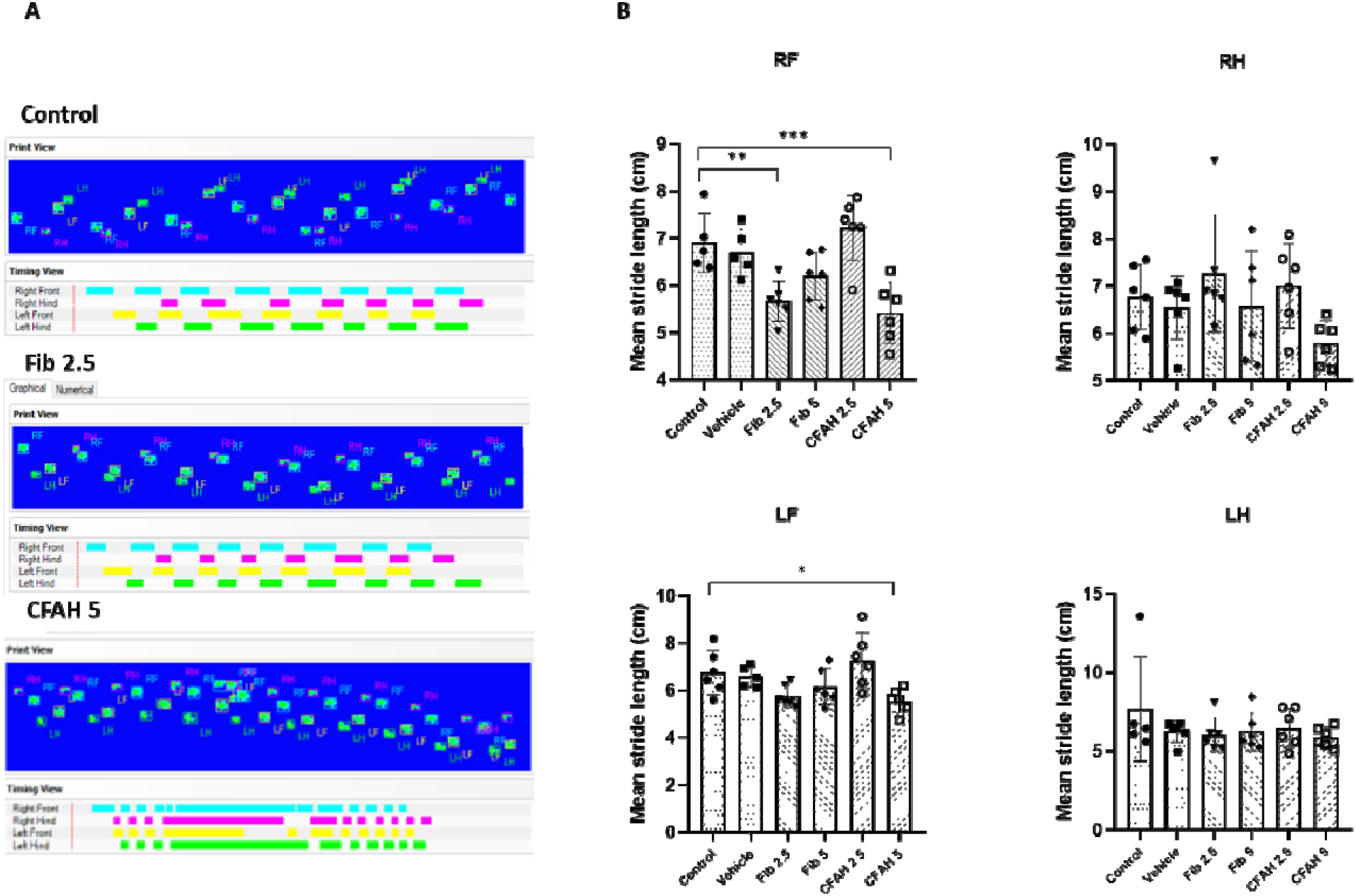
**A.** Representative pawprint impressions of control, Fib 2.5 and CFAH 5 on the catwalk gait analysis system. Note the shortening and drag in the prints of CFAH group. **B.** Histograms showing mean stride length of RF, RH, LF and LH. Note the significant reduction in the mean stride length of RF (control vs Fib 2.5 **p= 0.0058; control vs CFAH 5 ***p= 0.0009), and LF (control vs CFAH 5 *p= 0.0489) and no significant change in the stride length of RH & LH.

#### Effect on cognition

The fibrinogen-injected mice receiving 2.5µg dosage showed a mild decrease in discrimination index (DI), while that of CFAH-injected mice was significantly low (Figure 2A, vehicle vs CFAH5 p=0.0423). The percentage of exploration of the familiar object was significantly high in the CFAH-5 mice (Figure 2B, vehicle vs CFAH5 *p=0.0233). The percentage of exploration in other zones (non-exploration) along with overall exploration was significantly low in the CFAH -injected group (Figure 2D; vehicle vs CFAH 5 µg injected *p= 0.0428, Figure 2E; vehicle vs CFAH 5 µg injected *p= 0.0118).

**Figure 2.**
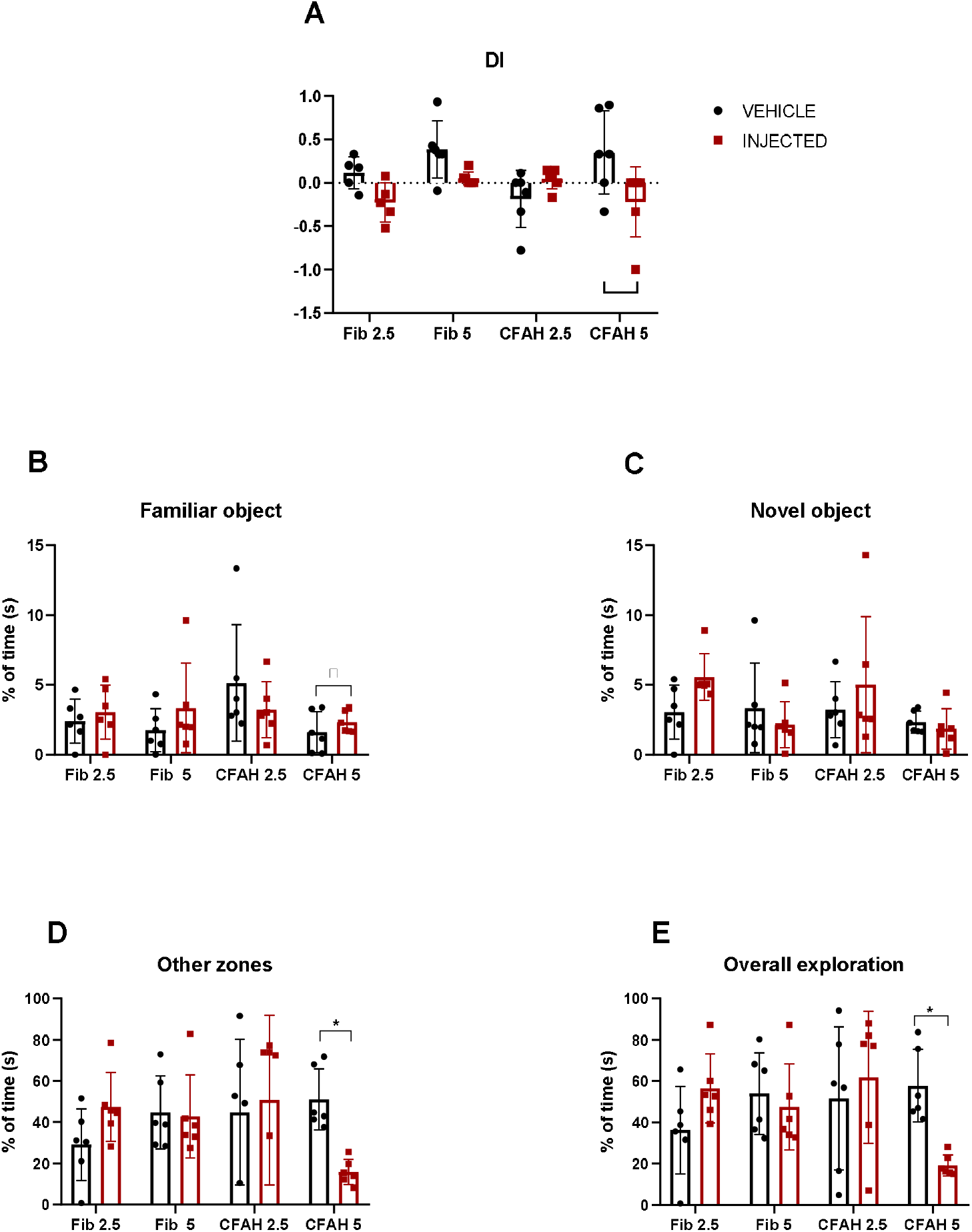
Histograms depicting different parameters of the NOR test with mean ± SD **A.** Note the trend of reduction in the discrimination index (DI) of Fib 2.5 and Fib 5 groups and a significant reduction in CFAH group (vehicle vs CFAH 5 p=0.0423) 5 **B.** Percentage of time spent with familiar objects where the CFAH 5 injected group shows a significantly high exploration time (vehicle vs CFAH 5, p=0.0423) and **C.** Novel objects **D.** Exploration of other zones (non-exploration) where the CFAH 5 group shows a significantly low exploration (vehicle vs CFAH 5 injected *p= 0.0428) **E.** In the overall exploration time, the CFAH 5 group shows a significantly low percentage (vehicle vs CFAH 5 injected *p= 0.0118).

#### Fibrinogen and CFAH cause depletion of DA neurons and induce cellular hypertrophy

The number of neurons, as quantified by the optical fractionator probe of unbiased stereology, showed a significant reduction in the total number of DA neurons (Figure 3F) in fibrinogen-injected mice (vehicle vs Fib 2.5 **p=0.0016; and vehicle vs Fib 5 **p=0.0063), but not in the CFAH injected ones. The number of degenerating DA neurons (Figure 3G) appeared to be relatively more in the CFAH-2.5, but the numbers did not reach statistical significance. The soma area was significantly larger in the higher dosage groups (vehicle vs Fib 5 µg **p=0.0046; vehicle vs CFAH 5 µg *p=0.0137) (Figure 3H), suggesting neuronal hypertrophy.

**Figure 3.**
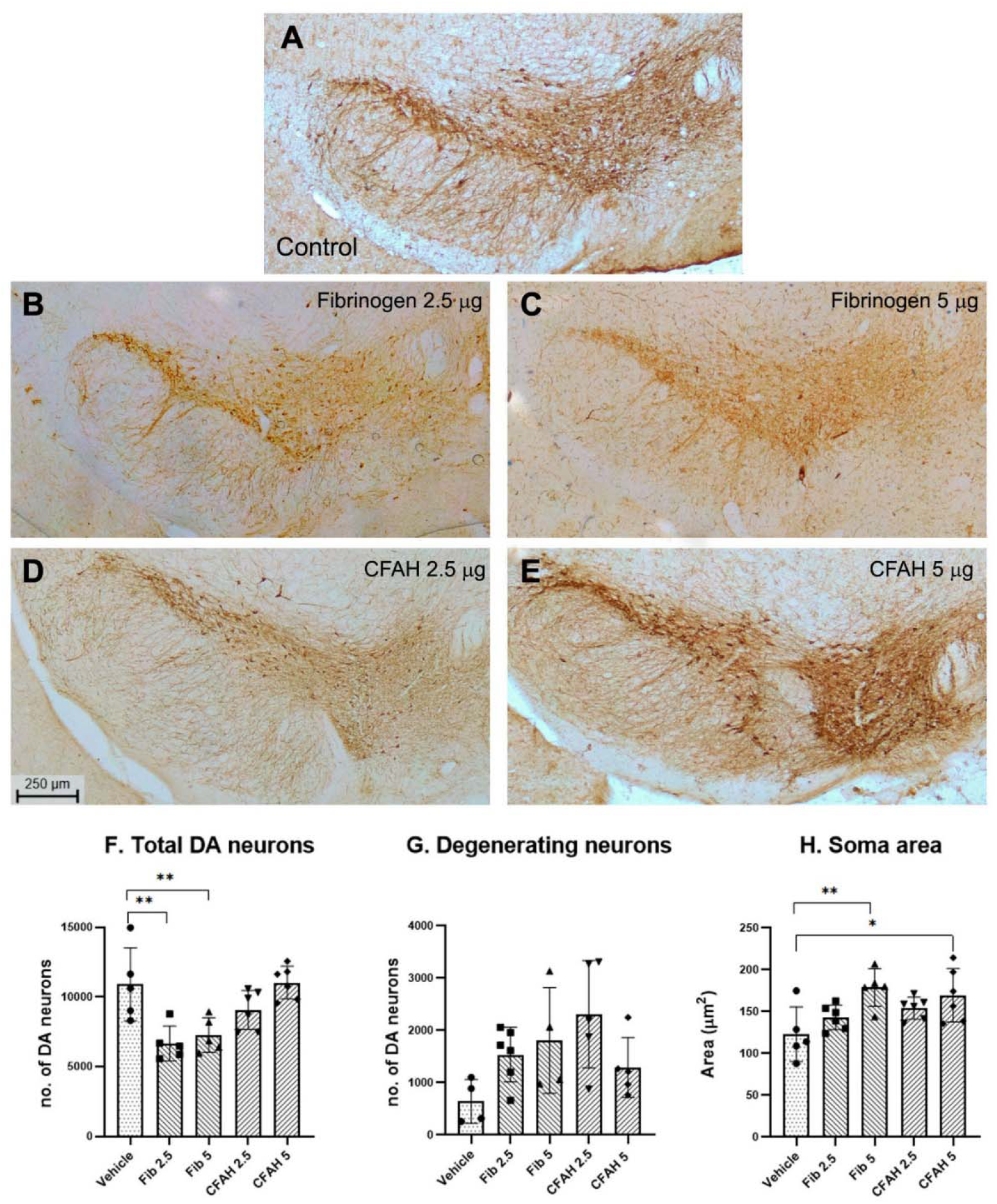
Representative low magnification photomicrographs of TH immunopositive DA neurons within SNpc of **A.** control **B.** Fib 2.5 **C.** Fib 5 **D.** CFAH 2.5 **E.** CFAH 5 **F.** The histogram shows a significant reduction in the number of DA neurons in Fib 2.5 and Fib 5 in comparison to vehicle (vehicle vs Fib 2.5 **p=0.0016), (control vs Fib 5 **p=0.0063) **G.** Note absence of significance in the number of degenerating DA neurons. **H.** A significant increase in the soma size of DA neurons in Fib 5 µg (vehicle vs Fib 5 **p=0.0046) and CFAH 5 (vehicle vs CFAH 5 *p=0.0137) mice was noted. Note the trend of increase in the soma size which measured to (16 %) in Fib 2.5 µg and (20 %) in CFAH 2.5 µg injected mice but did not reach significance. Scale bar = 250 µm for all micrographs.

#### Cytomorphometry based estimation of neuronal, cytoplasmic size and cellular TH levels

The neuronal clusters that form different sub-regions of the substantia nigra make topographical connections with specific sub-regions of the striatum. Hence, we segregated our observations accordingly in the VTA and SN. As regards the VTA, interesting observations confirm a significant reduction in the cytoplasmic TH in Fib 2.5 and Fib 5 (Figure 4F vehicle vs. Fib 2.5 *p=0.0244, vehicle vs Fib 5 *p=0.0307), which was complemented by cytoplasmic hypertrophy in Fib 5 **p=0.0024 (Figure 4G). The nigra was further sub-divided in medial, dorsal and lateral subdivisions. The DA neurons showed shrinkage of the cytoplasm in the fibrinogen 2.5 µg injected mice in the medial, dorsal and lateral aspects of the SN (Figure. 4N, 5G and 5N), while the expression of TH was unchanged (Figure 4M, 5F and 5M,).

**Figure 4:**
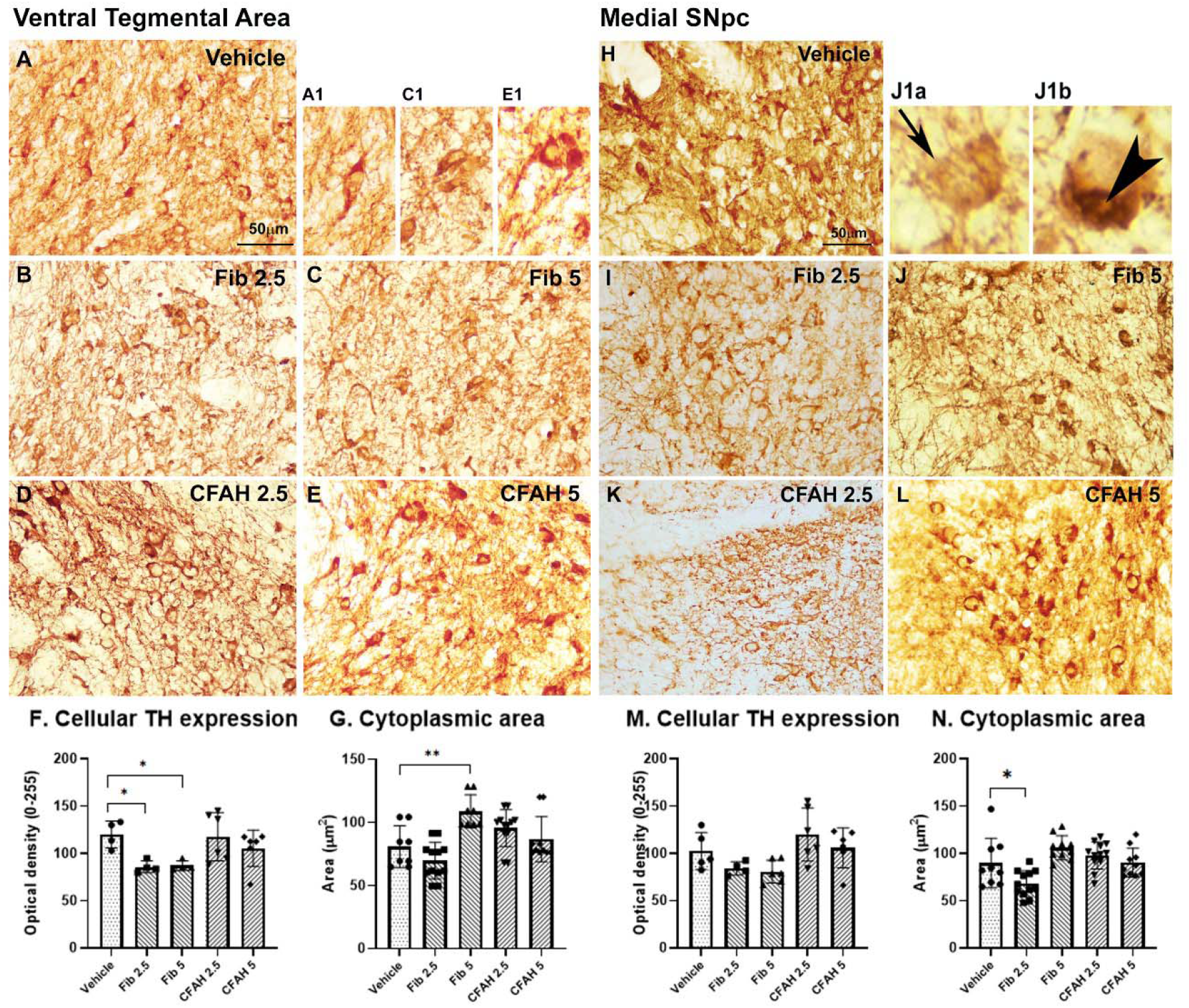
Representative photomicrographs of VTA DA neurons showing cellular TH expression in **A.** vehicle. **B.** Fib 2.5 **C.** Fib 5 **D.** CFAH 2.5 **E.** CFAH 5 administered study groups. **A1, C1, E1:** Photomicrographs showing VTA at 100X from control, Fib 5 and CFAH 5. **F.** Note the significant reduction in TH expression in Fib 2.5 and Fib 5 (vehicle vs. Fib 2.5 *p=0.0244, vehicle vs Fib 5 *p=0.0307) **G.** Significant increase in the cytoplasmic area (vehicle vs Fib5 **p=0.0024). Representative photomicrographs of nigral DA neurons showing cellular TH expression of the medial subregion in **H.** vehicle. **I.** Fib 2.5 **C.** Fib 5 **J1a.** Note the arrow pointing towards degenerating neurons and **J1b.** Arrowhead pointing towards a cytoplasmic vacuole of Fib 5. **K.** CFAH 2.5 **L.** CFAH 5. **M.** Medial SN showed no significant change in TH expression **N.** Note the significant decrease in the cytoplasmic area of Fib 2.5 (vehicle vs. Fib 2.5 *p=0.0327) to be noted. Scale bar =50 µm for all micrographs.

**Figure 5:**
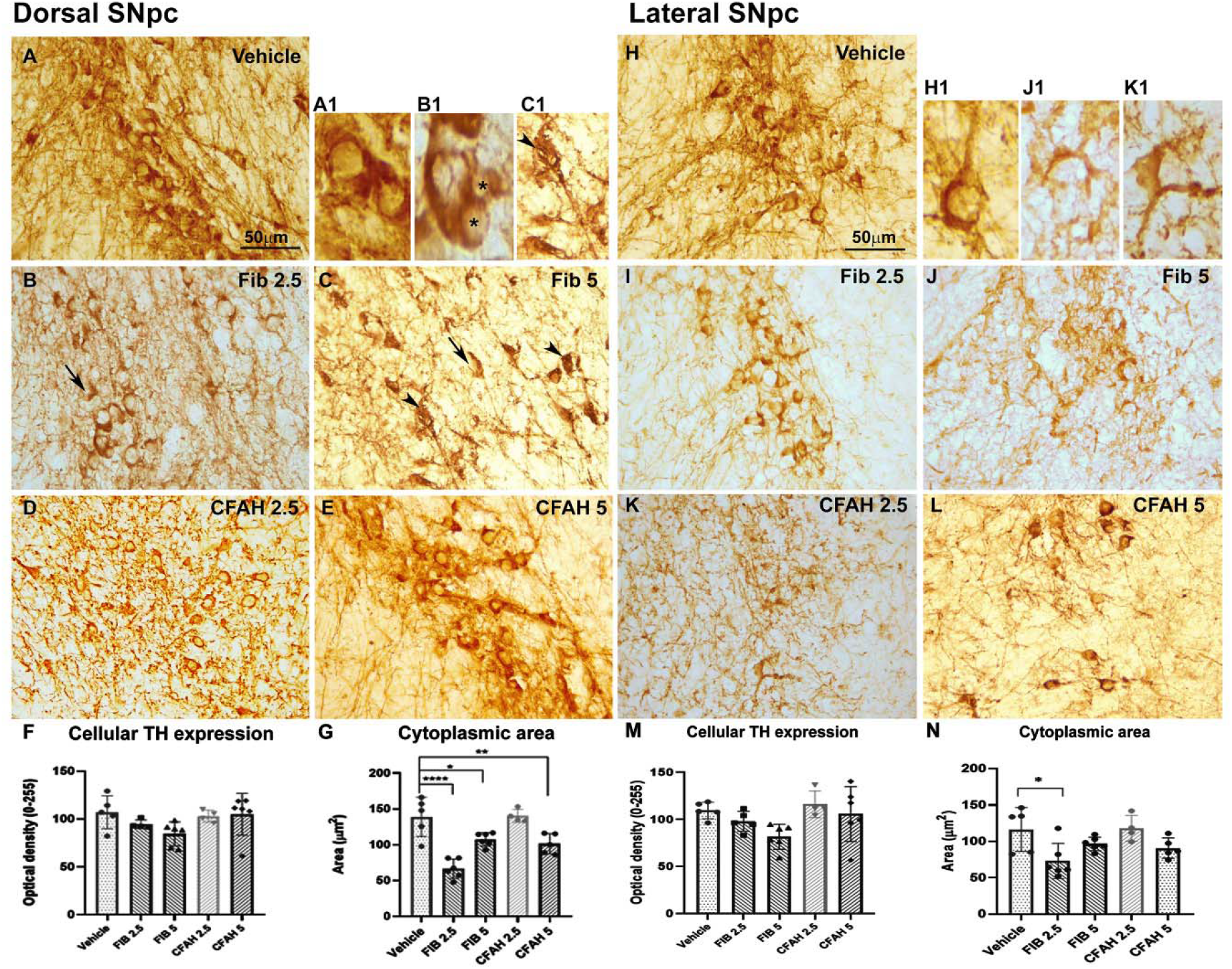
Representative photomicrographs of DA neurons of the dorsal (A-E, A1-C1) and lateral (H-L; H1-K1) subregions of the SNpc, showing pan-cytoplasmic TH expression. Note the alterations in the dorsal region (A-E, A1-C1) in vehicle injected mice **A.** Note the single healthy DA neuron at higher magnification (100X, A1). Note the neurons in fibrinogen and CFAH injected mice (**B-E). B.** Fib 2.5, **B1.** **represent smaller neurons **C.** Fib 5 **C1.** Note the degenerating neuron in Fib 5 (arrow-heads). **D.** CFAH 2.5 **E.** CFAH 5. **F.** Dorsal SN showed no significant changes in TH expression. **G.** Note the significant decrease/shrinkage in dorsal cytoplasmic area (vehicle vs Fib 2.5 ****p=0.0002). Representative photomicrographs of the DA neurons of the lateral subregion (H-L) of the SNpc showing cellular TH expression in **H.** vehicle. **H1.** Note the single healthy DA neuron at higher magnification (100X) **I.** Fib 2.5 **J.** Fib 5, **K.** CFAH 2.5 **J1**. Photomicrographs at higher magnification (100x) show arrows pointing towards the degenerating DA neuron in Fib 5 and **K1** CFAH 2.5 **L.** CFAH 5. **M.** Histogram showing the trend of decrease in TH expression in lateral SN in both Fib 2.5 (10 %) and Fib 5 (26 %) groups. **N.** Also note the shrinkage in the cytoplasmic area (vehicle vs Fib 2.5 *p=0.0061). Scale bar =50 µm for all micrographs.

#### Effects of fibrinogen and CFAH on the striatal TH expression

The striatum is the main input center of the basal ganglia, which is innervated by the mesencephalic DA neurons. Complementing the loss of DA neurons of the SNpc, a significant reduction was noted in the TH expression in the striatum (Figure 6P) (vehicle vs Fib 2.5 *p=0.0429; vehicle vs Fib 5 **p=0.0039) and dorsolateral striatum (Figure 6Q) (vehicle vs Fib 5 **p=0.0083). However, no alterations were observed in the ventral parts of the striatum consisting of the nucleus accumbens. The expression in the CFAH groups also remained unaltered.

**Figure 6:**
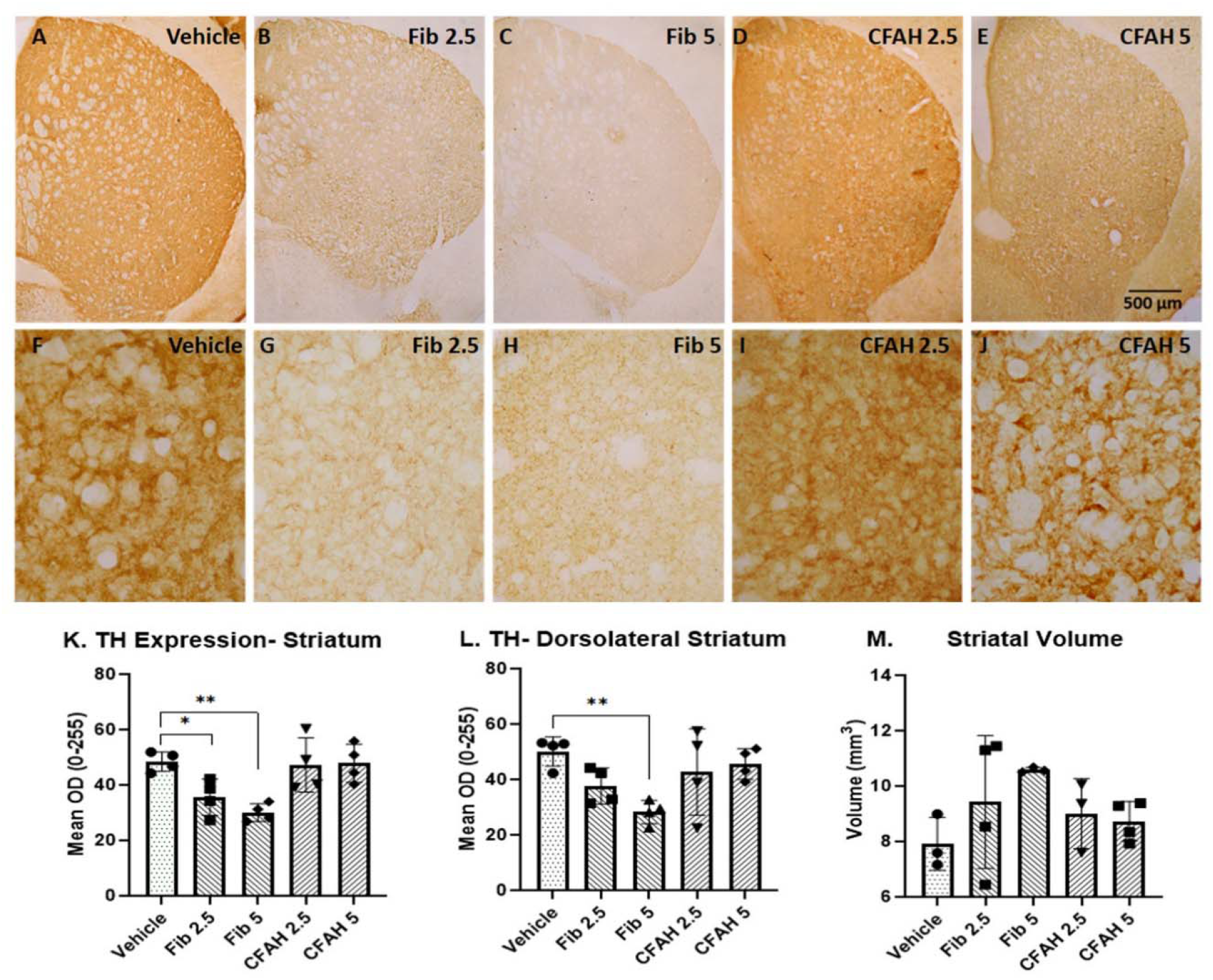
Representative photomicrographs of striatum (overall striatum) showing TH expression in the terminals across various groups. The upper panel shows low magnification (4X) images of different study groups i.e. **A.** Vehicle **B.** Fib 2.5 **C.** Fib 5 **D.** CFAH 2.5 **E.** CFAH 5. The lower panel shows high magnification (40X) images of the dorsolateral striatum (DL) different study groups i.e. **F.** Vehicle **G.** Fib 2.5 **H.** Fib 5 **I.** CFAH 2.5 **J.** CFAH 5 **K.** Histograms showing differential expression of TH in terms of OD in the overall striatum (Vehicle vs Fib 2.5 *p=0.0429; Vehicle vs Fib 5 **p=0.0039). Note the significant decrease in the overall striatal TH expression of Fib 2.5 and Fib 5 groups. and **L.** dorsolateral striatum (DL) shows a decrease in the expression (Vehicle vs Fib 5 **p=0.0083) **M.** No significant changes are seen in striatal volume. Scale bar of all micrographs =500 µm

#### Neurons Hypertrophy in the hippocampal subfields

Hippocampal subfield atrophy has implications in cognitive impairment, besides, DA neurons also innervate the hippocampal subfields like CA-1 and subiculum. Presence of hypertrophic nucleus and cytoplasm of CA-1 neurons was observed in fibrinogen and CFAH-injected groups (Figure 7F, vehicle vs Fib 2.5 **p= 0.0079, vehicle vs Fib 5 **p= 0.0006, vehicle vs CFAH 5 µg *p=0.163; Figure 7G, vehicle vs Fib 2.5 **p= 0.0083, vehicle vs Fib 5**p= 0.0096, vehicle vs CFAH 5 *p=0.0384) respectively. Similarly, the nucleus of the subicular cells appeared enlarged, in response to both fibrinogen and CFAH-injection (Figure 7N, vehicle vs Fib 2.5 µg **p=0.0017, vehicle vs Fib 5 µg ***p= 0.0003, vehicle vs CFAH 5 µg **p=0.0085) whereas cytoplasmic hypertrophy was noticed only in fibrinogen-injected groups (Figure 7O, vehicle vs Fib 2.5 **p=0.0025, vehicle vs Fib 5 *p= 0.0064). No overall volumetric changes were observed in the CA1 and subiculum (Figure 7H and 7P).

**Figure 7:**
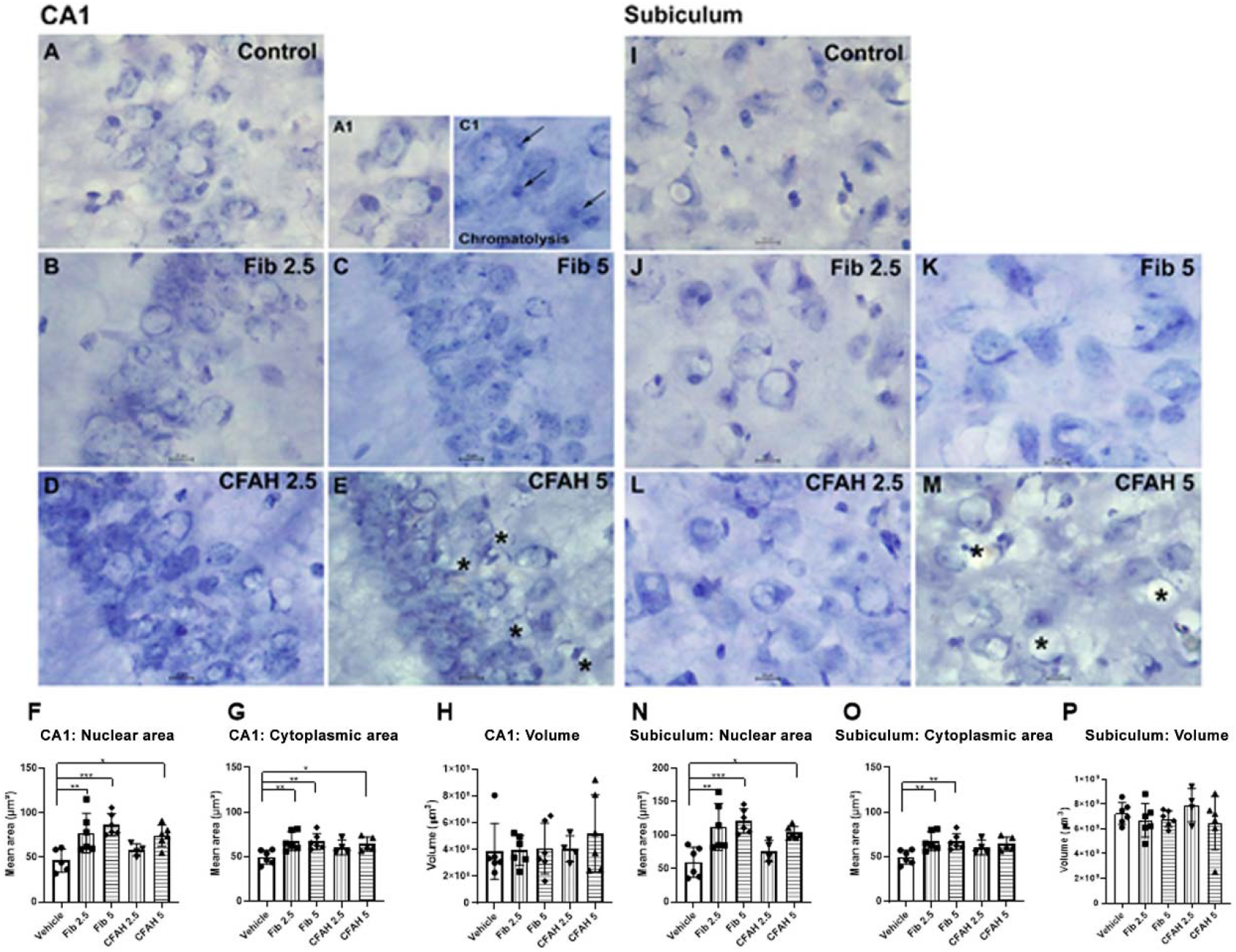
Representative Low magnification photomicrographs of mice CA 1 following different injection regimes for the proteins i.e., fibrinogen and CFAH **A.** vehicle **B.** Fib 2.5 **C.** Fib 5 **C1:** Arrows indicate chromatolysis, **D.** CFAH 2.5 **E.** CFAH 5 **E1**: **** (asterisks) represent necroptotic cells. Histograms showing the morphometric index of CA1 neurons **F.** Note the significant increase in the area of the neuronal nucleus of Fib 2.5 µg, Fib 5 µg, CFAH 5 µg injected mice (vehicle vs Fib 2.5 µg **p= 0.0079, vehicle vs Fib 5 µg **p= 0.0006, vehicle vs CFAH 5 µg *p=0.163), and **G.** a significant increase in the cytoplasmic area of the neurons (vehicle vs Fib 2.5 **p= 0.0083, vehicle vs Fib 5**p= 0.0096, vehicle vs CFAH 5 µg *p=0.0384). **H.** Histogram showing unaltered volume of the CA1. **I.** vehicle **J.** Fib 2.5 **K.** Fib 5 **L.** CFAH 2.5 injected animals showed patches of hypertrophied cells **M.** Note the ‘necroptotic spots’ (**asterisks) in the subiculum of CFAH 5 study group, indicating degenerative changes. **N.** The histograms confirm nuclear enlargement of the neuronal nucleus of Fib 2.5µg, Fib 5µg, CFAH 5µg (vehicle vs Fib 2.5 µg **p=0.0017, vehicle vs Fib 5 µg ***p= 0.0003, vehicle vs CFAH 5 µg **p=0.0085), **O.** as well as cytoplasmic enlargement. Note the significant increase in the cytoplasmic area (vehicle vs Fib 2.5 **p=0.0025, vehicle vs Fib 5 **p= 0.0064) in Fib 2.5 and Fib 5 study groups, both suggesting cellular hypertrophy. **P.** Histogram showing unaltered volume. Scale bar =10 µm.

**Figure 8:**
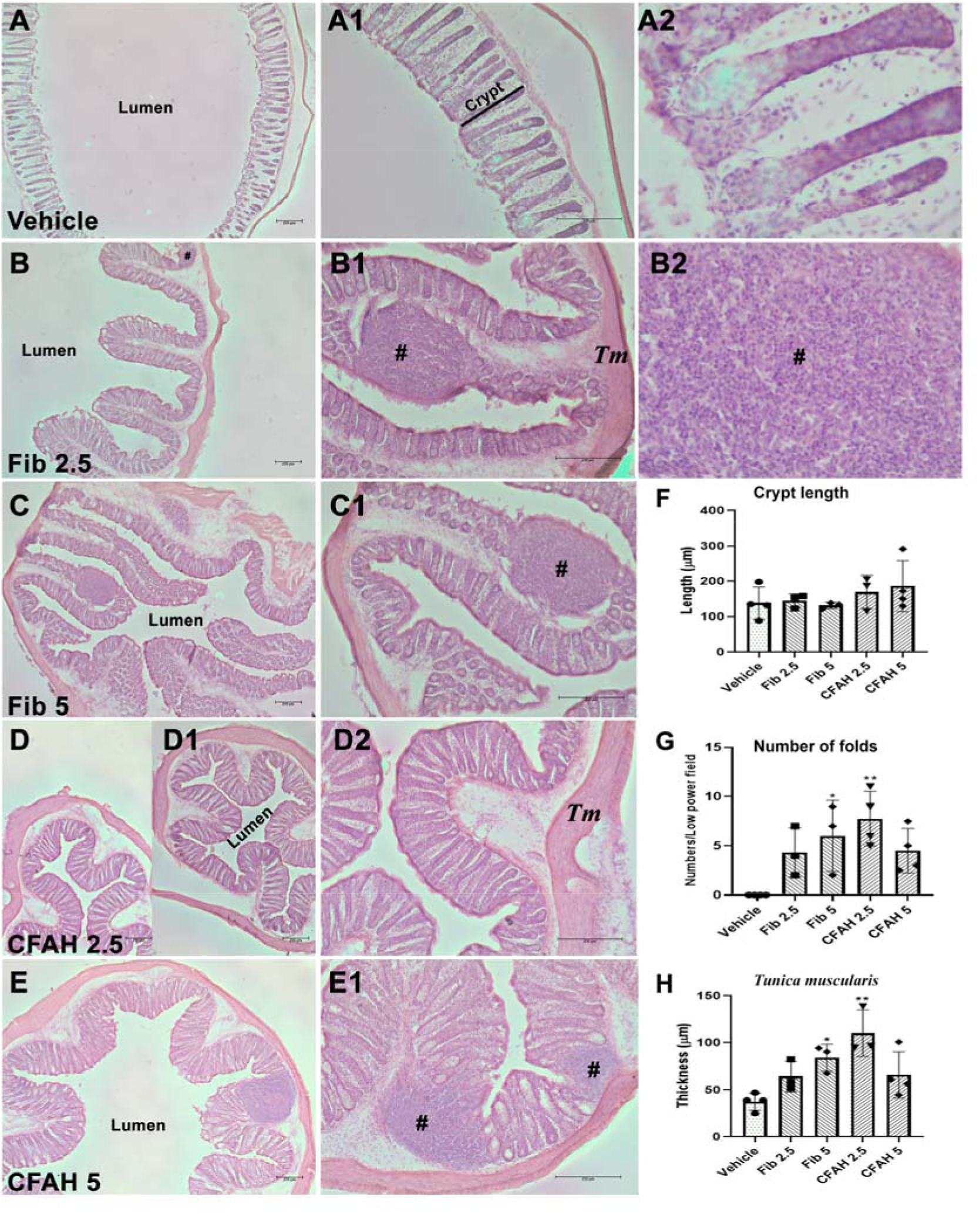
Representative Low magnification photomicrographs of mice colon under control (vehicle) conditions (A-A2) and following different injection regimes for fibrinogen (B-B2 and C-C1) and CFAH (D-D2 and E-E1) at different magnifications (4X and 10X). **B.** Fib 2.5 **C.** Fib 5 C1. Note the presence of inflammatory patches (##; B-E1) in the guts of mice injected with either Fib or CFAH. Note the presence of significantly higher number of folds in colonic lumen and the considerable thickening of the Tunica muscularis in the injected groups (*control versus fib 5 p< 0.05; **control versus CFAH 2.5 p< 0.01).

#### Alterations in colonic architecture

The colon showed few alterations in response to both fibrinogen and CFAH. The colonic crypt lengths, were preserved between groups, although the average length was marginally higher in the CFAH 5 group. The controls showed no folds, although wavy mucosa was noted. Interestingly, the folds were higher in the experimental groups viz. Fib 5 (*p <0.05) and CFAH 2.5 (**p <0.01), vis-à-vis the controls. The folds were irregular sized, long, narrow, compact, bulgy, broken, disintegrated, large, rounded, etc. The increase in the number of folds effectively reduced the size of the colonic lumen, which may possibly reduce the peristaltic movement and results in constipation; which is a prodromal symptom of PD. Another interesting observation was the thickening of the tunica muscularis in the injected groups i.e. Fib 5 (*p <0.05) and CFAH 2.5 (**p <0.01), vis-à-vis the controls. Presence of Peyer’s-like colitic patches (##) in the injected groups suggest an inflammatory response.

## Discussion

Studies to understand the neurobiology of PD pathogenesis have often been restricted to the use of neurotoxins and transgenic mice, including those over expressing α-synuclein (Magen et al., 2012; Elabi et al., 2021). In the present study we examined the role of endogenous molecule fibrinogen, following exogenous supplementation; in causing or aggravating PD pathology. Although fibrinogen-induced BBB breach has been implicated in other neurodegenerative disorders like Alzheimer’s disease, occasional studies have focused on its application in generating animal models of PD. A recent study demonstrating BBB breach in α-synuclein transgenics and the over expression of fibrinogen in the mice striatum, opened up the possible bi-directionality of the pathological events, wherein fibrinogen may also be capable of inducing α-synuclein associated deficits. Our study provides novel insights into the pathogenic potential of the proteins fibrinogen and CFAH in terms of neurobehavioral and neuroanatomical correlates associated with PD and cognitive impairment.

### Both Fibrinogen and CFAH induce behavioral deficits in mice

Shortening of the stride length in fibrinogen and CFAH injected mice provides the first evidence of shuffling steps, a cardinal clinical manifestation noted in PD patients (Morris et al., 1996). As evident from the alterations in paw prints, there was a drag in the movement of CFAH-injected mice that can be corroborated well with that reported in PD patients (Kim et al., 2018). Walking is not solely a motor task; it necessitates cognitive maneuvering for the coordination of movement and maintaining postural stability. Various studies, in humans, have reported that short stepped gait is an additional predictor of cognitive decline (Savica et al., 2017). Consequentially, we attempted to examine the cognitive implications of higher levels of fibrinogen and CFAH.

Heretofore, studies have shown recognition memory deficits, as assessed by NOR test, in three distinct models, i.e., in an α-synuclein overexpressing mouse model (Magen et al., 2012); in mice infused with α-synuclein oligomers within brain (Fortuna et al. 2017) and in mice receiving intraperitoneal injections of MPTP (Moriguchi et al., 2012). The trend of low discrimination index in our fibrinogen (2.5µg) and CFAH (5µg) injected mice, substantiates the induction of cognitive deficits by these proteins. The enhanced exploration in the familiar object zone by the fibrinogen injected mice (2.5µg) hints at compromised exploration or loss of investigatory preferences of the novel object in them whereas the severely reduced ability of CFAH (5µg) injected mice in exploring the entire arena, might be a reflection of larger influence of CFAH on the motor system than the cognitive component as also seen in our gait experiments. Further, we cannot preclude the possible loss of motivation, another non-motor symptom of PD, a significant feature of PDCI, as a reason for the reduced exploration following CFAH (5µg) administration.

### Fibrinogen induces neuroanatomical alteration in the SNpc and Striatum

A massive loss of TH immunoreactive DA neurons in the SNpc is the hallmark neuropathological characteristic of PD (Hirsch et al., 1988) The approximate loss of 36% DA neurons in the SNpc of fibrinogen-injected mice compared to the controls, reflects the ability of fibrinogen to cause frank degeneration of DA neurons, which may form the basis for motor deficits. The toxicity of fibrinogen, in our study, was moderate, compared to the >50% reduction in the DA neuronal numbers earlier reported in the substantia nigra of MPTP injected mice (Vidyadhara et al. 2017) or in patient brains (Hirsch et al., 1988). However, in situations simulating disease condition i.e., where high levels of fibrinogen may be in continuous circulation, it is likely to enhance the magnitude of loss. Our results also partly concur with the findings of Alam et al., (2017), who showed that a single injection of 20mg/kg MPTP through i.p. route causes a loss of ∼60% of TH immunoreactive neurons in the SN. Although CFAH failed to induce active neuronal loss in the mice, the number of degenerative neurons were fairly high (CFAH 2.5; ∼16 %), hinting at an active phase of degeneration, and a need for long term follow-up.

An overall hypertrophy of DA neurons in the medial, dorsal and lateral subfields of SNpc observed in our study could be a compensatory mechanism against the DA cell loss in the SNpc. It resembles the age-related hypertrophy of cell bodies of the DA neurons of the SNpc of humans (Cabello et al., 2002), and vervet monkeys (Janson et al., 1991). Apoptosis is an important phenomenon in dying DA neurons of PD patients and cellular shrinkage is a vital morphological feature of apoptosis (Anglade et al., 1997; Vidyadhara et al., 2017) and hence the reduction of the cytoplasmic area noted in the DA neurons of the medial, dorsal, and lateral aspects of the SNpc in the fibrinogen injected mice appear to be a prelude to apoptosis which also corroborates with our recent findings demonstrating increases in apoptotic factors in MPTP-injected C57BL/6J mice with lower number of DA neurons (Yarreiphang et al., 2023). The overall hypertrophy of neurons accompanied by cytoplasmic shrinkage are distinct anomalies that are seemingly contradictory and difficult to be explained.

In patients with dementia, lesions in the VTA caused dysexecutive syndrome (Adair et al., 1996). Hence, the hypertrophy of the VTA-DA neurons observed in our study may be an indirect effect of fibrinogen, on cognition. Age associated degenerative changes in the substantia nigra were earlier identified by the reduction in the TH expression within these structures, although the cell bodies were preserved; thus symbolizing “loss of phenotype” of the DA neurons (Chu & Kordower, 2007). The nigral DA neurons of the Caucasians who are inherently more susceptible to PD showed such a loss (Chu & Kordower, 2007); which was proposed to be a cellular index of susceptibility, whereas A9 neurons in Asian midbrains retained TH expression through aging. Asian Indians are less susceptible to PD compared to the Caucasians and hence, such preservation of phenotype may be an important feature of resilience (Alladi et al., 2009; Das et al., 2010; Pitcher et al., 2018, Je et al., 2021). Some neuroanatomical bases, pertaining to preservation of substantia nigra, have earlier been investigated (Alladi et al., 2009; Alladi et al., 2010a; Alladi et al., 2010b; Naskar et al., 2019; Jyothi et al., 2015). However, the prevalence rate of cognitive deficits in PD (PDCI), based on ethnicities, is yet not ascertained (Willis et al., 2012).

The striatum, the main input center of the basal ganglia, is innervated by DA projections from the substantia nigra and the VTA, forming the nigrostriatal connections. The loss of TH in the entire striatum suggests the expanse of the loss of DA inputs and partly corroborates the findings in MPTP-injected C57BL/6J mice (Vidyadhara et al., 2017). Although TH expression was fairly affected in the SNpc of the injected mice, the significantly low expression of TH in the VTA of fibrinogen injected mice indirectly indicates the effect of fibrinogen on cognition. Unlike the former observation there was no volumetric shrinkage which could be because of the shorter period of exposure, i.e., 48 hours.

Similar to the DA neurons of certain sub-regions of SN-VTA, a compensatory hypertrophy was also observed in the nucleus and cytoplasm of the CA1 and subicular cells. Although no volumetric shrinkage in CA1 and subiculum was noticed in the protein-injected groups. Enhancement of the DNA and RNA synthesis by the surviving neurons provides a plausible explanation for the enlargement of nucleus and cytoplasm to balance the ongoing or anticipated loss of neurons (Iacono et al., 2008). Several groups have earlier reported that CA1 (Ziehn et al., 2010; Mueller et al., 2010), CA2 (Mattila et al., 1999), CA3 (Wisse et al., 2014; Lenka et al., 2018) and the subiculum are more prone to degeneration in normal aging and neurodegenerative disorders, associated with cognitive impairment. Therefore, atrophy of the cells in hippocampal subfields may theorize cognitive decline (Iacono et al. 2008, Foo at al., 2016). MRI-based volumetric studies showed, atrophy of the hippocampal subfield as an inkling of cognitive impairment in PD, although PD patients with hippocampal atrophy may not display cognitive impairment (Camicioli et al., 2003).

The presence of irregular sized, long, narrow, compact, bulgy, broken, disintegrated, large or rounded colonic folds suggest that both the proteins bear the potential to induce micro-architectural changes in the gut. The increase in folds effectively reduced the colonic lumen size, which may affect the peristaltic movement and possibly results in constipation, an important prodromal symptom of PD. Another interesting observation was the thickening of the tunica muscularis in the injected groups. Fibrinogen is expressed endogenously to maintain the epithelial cells of the colon, therefore, I.P. injection of fibrinogen may enhance the process of replenishment of the epithelial layer, resulting in thickening of the Tunica muscularis (Seltana et al., 2022). Presence of Peyer’s-like colitic patches (##) in the injected groups suggest an inflammatory response.

It was long believed that fibrinogen, the plasma protein, can enter the brain only if there is a breach of the BBB and induce inflammatory changes (Davalos & Akassoglou 2012). However, using *vitro* modalities, Tyagi et al (2008) suggested that even raised levels of fibrinogen enhances endothelial cell permeability through the extracellular signal regulated kinase particularly in cardiovascular and cerebrovascular diseases. Elevated levels of fibrinogen and CFAH in the peritoneum of mice, following injections, mimic clinical conditions of increased CSF levels in PDCI patients. The PDCI like neurobehavioral and neurodegenerative changes in the C57BL/ 6J mice brain, tempt us to hypothesize that the elevated levels of these proteins are capable of affecting the BBB by binding to their cognate receptors, which is trailed by the neurodegeneration. The increase in folds within the colon effectively reduced the colonic lumen size, which may result in reduced peristaltic movement and possibly in constipation; is a prodromal symptom of PD.

Reduced expression of complement component 3 (C3) is implicated in several bowel diseases viz. the Crohn’s disease, inflammatory bowel disease, and ulcerative colitis, as studied using knock-out modality (Park et al., 2020 preprint). CFAH being an active member of the complement system; a valid correlation may be speculated between enhanced levels of CFAH and constriction of lumen in the CFAH injected mice. It is, thus tempting to surmise that CFAH also affects the gut and the innate immune system.

The frank loss of neurons following fibrinogen injection and the observation of active degenerative profiles upon CFAH administration, suggest that when present in combination, as in patients’ CSF, they may cross-complement the neurotoxicity. This also points at the need to investigate other aspects of neurodegeneration. The behavioral deficits provide the second objective evidence of their pathogenic potential and support the vital clinical symptoms of PD. Taken together, our observations suggest that fibrinogen being present in comparatively higher levels than CFAH, it bears a slightly higher potential for inducing pathogenesis.

## Conclusion

Our study is the first of its kind that offers an animal model for understanding the etiopathogenesis of PDCI. It also provides valuable assistance to validate the pathogenicity of the identified markers in causing PDCI like symptoms. Based on the neurobehavioral and neuropathological deficits that are induced by fibrinogen and CFAH, it may be surmised that these two proteins are putative biomarkers of PDCI. In addition, their increased levels in the CSF might be indicative of the BBB breach that leads to neurodegenerative changes associated with PD and further ascertainment on autopsied patient tissues may provide vital leads. In the given experimental conditions, fibrinogen proves to be more potent in causing PDCI like neurobehavioral and neuropathological changes, thus its significantly higher levels in CSF may align well with the severity of motor and cognitive pathology in the PDCI patients. Further investigations are required to assess their combinatorial effects.

## Abbreviations

BBB: blood-brain barrier
CFAH: complement factor H
CI: cognitive impairment
Fib: fibrinogen
NOR: novel object recognition
PD: Parkinson’s disease
SNpc: substantia nigra pars compacta
VTA: ventral tegmental area

## Author contributions

AN performed the experiments, analyzed the data and wrote the first draft of the manuscript. S.M. performed the quantification of the nigral neurons using densitometry. S.S. quantified the intensity of striatal TH and the hippocampal cytomorphology. PB performed the quantification of gut histology. P.A.A., conceptualized the study, designed the experiments, captured images and analyzed the data. P.A.A., S.H, R.Y. and P.K.P. obtained funds and gave critical suggestions. P.A.A. revised the manuscript and prepared the final article. All the authors have approved the final version of the MS.

## Acknowledgements

The authors are grateful to the Indian Council of Medical Research, for funding support to P.A.A. (BMS/TF/Trans-Neuro/2014-3424/Dec-15/48/ MH/Govt). A.N. was an ICMR-JRF [no. 3/1/2/JRF-2014/HRD-102 (40832)]. The authors sincerely acknowledge the help of Dr. Sumana Chakravarty for her suggestions in designing the behavioral studies and Dr. Anita Mahadevan and Ms. Shakti for facilitating hematoxylin staining in the human brain tissue repository (HBTR), NIMHANS.

